# Need for high-resolution Genetic Analysis in iPSC: Results and Lessons from the ForIPS Consortium Authors

**DOI:** 10.1101/379818

**Authors:** Bernt Popp, Mandy Krumbiegel, Janina Grosch, Annika Sommer, Steffen Uebe, Zacharias Kohl, Sonja Plötz, Michaela Farrell, Udo Trautmann, Cornelia Kraus, Arif B. Ekici, Reza Asadollahi, Martin Regensburger, Katharina Günther, Anita Rauch, Frank Edenhofer, Jürgen Winkler, Beate Winner, André Reis

**Affiliations:** Institute of Human Genetics, University Hospital Erlangen, Friedrich-Alexander-Universität Erlangen-Nürnberg (FAU), Schwabachanlage 10, 91054 Erlangen, Germany; Department of Molecular Neurology, University Hospital Erlangen, Friedrich-Alexander-Universität Erlangen-Nürnberg (FAU), Schwabachanlage 6, Erlangen, Germany; Department of Stem Cell Biology, University Hospital Erlangen, Friedrich-Alexander-Universität Erlangen-Nürnberg (FAU), Glückstrasse 6, Erlangen, Germany; Stem Cell Biology and Regenerative Medicine Group, Institute of Anatomy and Cell Biology, Julius-Maximilians-University of Würzburg, Würzburg, Germany; Institute of Medical Genetics, University of Zurich, Schlieren-Zurich, Switzerland

**Author notes:** Co-first author (Bernt Popp, Mandy Krumbiegel). Corresponding author (A.Re.). **Competing Interests** The authors declare no competing interests.

**Keywords:** iPSC, ForIPS, genetic quality control, exome sequencing, chromosomal microarray

## Abstract

Genetic integrity of induced pluripotent stem cells (iPSCs) is essential for their validity as disease models and for potential therapeutic use. We describe the comprehensive analysis in the ForIPS consortium: an iPSC collection from donors with neurological diseases and healthy controls. Characterization included pluripotency confirmation, fingerprinting, conventional and molecular karyotyping in all lines. In the majority, somatic copy number variants (CNVs) were identified. A subset with available matched donor DNA was selected for comparative exome sequencing. We identified single nucleotide variants (SNVs) at different allelic frequencies in each clone with high variability in mutational load. Low frequencies of variants in parental fibroblasts highlight the importance of germline samples. Somatic variant number was independent from reprogramming, cell type and passage. Comparison with disease genes and prediction scores suggest biological relevance for some variants. We show that high-throughput sequencing has value beyond SNV detection and the requirement to individually evaluate each clone.

## INTRODUCTION

Genetic variants influence cellular mechanisms, thus leading to specific phenotypic presentations in the organism, both in rare and common disease. Neurological disorders like Parkinson’s disease (PD) typically comprise both rare and common genetic risk variants with large and small effect sizes, respectively. Studying the pathomechanism in patient cells is often limited because the disease relevant tissues are not accessible. Human embryonic stem cells (ESC) can be differentiated into cells from all three germ layers (endoderm, mesoderm, ectoderm) but pose legal and ethical issues. In contrast, induced pluripotent stem cells (iPSCs) can be derived from adult tissues using exogenous expression of four transcription factors (*POU5F1*, *SOX2*, *KLF4*, *MYC*) and can be differentiated into somatic cells *in vitro*.^1–3^ Human iPSCs promise not only easy access to cells for scientists interested in disease modelling but also personalized medicine for patients affected by rare diseases.

While different protocols (non- / integrating viral, non-integrating non- / viral) for the generation of iPSC lines have been established, quality control (QC) during reprogramming, differentiation and culturing steps remains an area of active development.^4^ Loss of genetic integrity as a source of variability in iPSCs^5^ and in therefrom derived cells is a possible confounder compromising their validity as disease models. Certain genetic variants could be associated with increased risk of cancer or dysfunction when using these cells for regenerative therapeutic interventions. Indeed, tumorigenicity has been reported in transplanted stem cells,^6^ and a recently published clinical trial using autologous iPSC derived retinal cells^7^ was temporarily halted due to concerns of tumorigenic potential. Finally, somatic *TP53* mutations previously identified in tumors were found in iPSC lines by applying exome sequencing.^8^ Taken together, a detailed characterization of genetic differences between donor and derived cells should be a central part of any iPSC-QC pipeline to ensure validity and safety.

Several groups and large consortia have studied the origin, quality and quantity of genetic variants found in iPSCs but absent from the donor’s germline.^5,9–11^ There is high variability in the methods used and the results reported. Also, the nomenclature for variants of different origin is inconsistent and often derives from research on cancer and developmental disorders.

Aneuploidies affecting the number of whole chromosomes in a cell are widely accepted as undesirable aberrations with potentially large effects in cells. Hence, conventional karyotyping is a standard QC measure used to detect these abnormalities in iPSCs. Similarly, somatic copy number variants (CNVs) like microdeletions and -duplications, typically comprising several genes or regulatory elements, are unfavorable. Although CNVs can be detected using chromosomal microarrays (CMA), this technique is not yet generally used to investigate iPSCs. High-throughput sequencing methods (“next-generation sequencing”; NGS) have enabled the exome and genome wide detection of single nucleotide variants (SNVs/indels). Several reports have shown considerable load of SNVs in iPSC.^12–15^

Here, we describe the ForIPS stem cell biobank resource, a national consortium with the primary goal to establish iPSC technologies to study molecular and cellular mechanisms involved in neurological disorders like PD. We present our approach to a stringent genetic workup, including conventional karyotyping, genetic fingerprinting and CMA in all cell samples. We report results of high coverage exome sequencing in a subset of this cohort selected to establish a suitable pipeline for iPSCs.

## RESULTS

### Characteristics of individuals included and iPSCs generated in the ForIPS biobank resource

The ForIPS study (Figure 1A) included 23 individuals (11 females and 12 males) of which 9 individuals (5 females, 4 males) were healthy controls without any neurologic disease (CT), 14 were patients affected (AP) by one of three neurological diseases: PD (1 female, 8 males), hereditary spastic paraplegia (HSP, *SPG11* gene, OMIM #604360 and *610844; 3 females), monogenic intellectual disability (ID; 2 females). The age at donation of fibroblasts ranged from 22 to 73 years (y) with a median of 45y. In CTs the age range was 23 to 70y with a median of 45y, and in APs the age range was 22 to 73y with a median of 45.5y. The oldest subgroup included individuals with PD (age range 36 to 73y, median 54y). Nine individuals were members of 4 families: “J2C” and “JF” are father and son, “88H”, “O3H” and “82A” are siblings, “PT1” and “CT1” are siblings and “55O” and “G7G” also are siblings (Figure 1B; see also Figure S1 and File S1).

**Figure 1.**
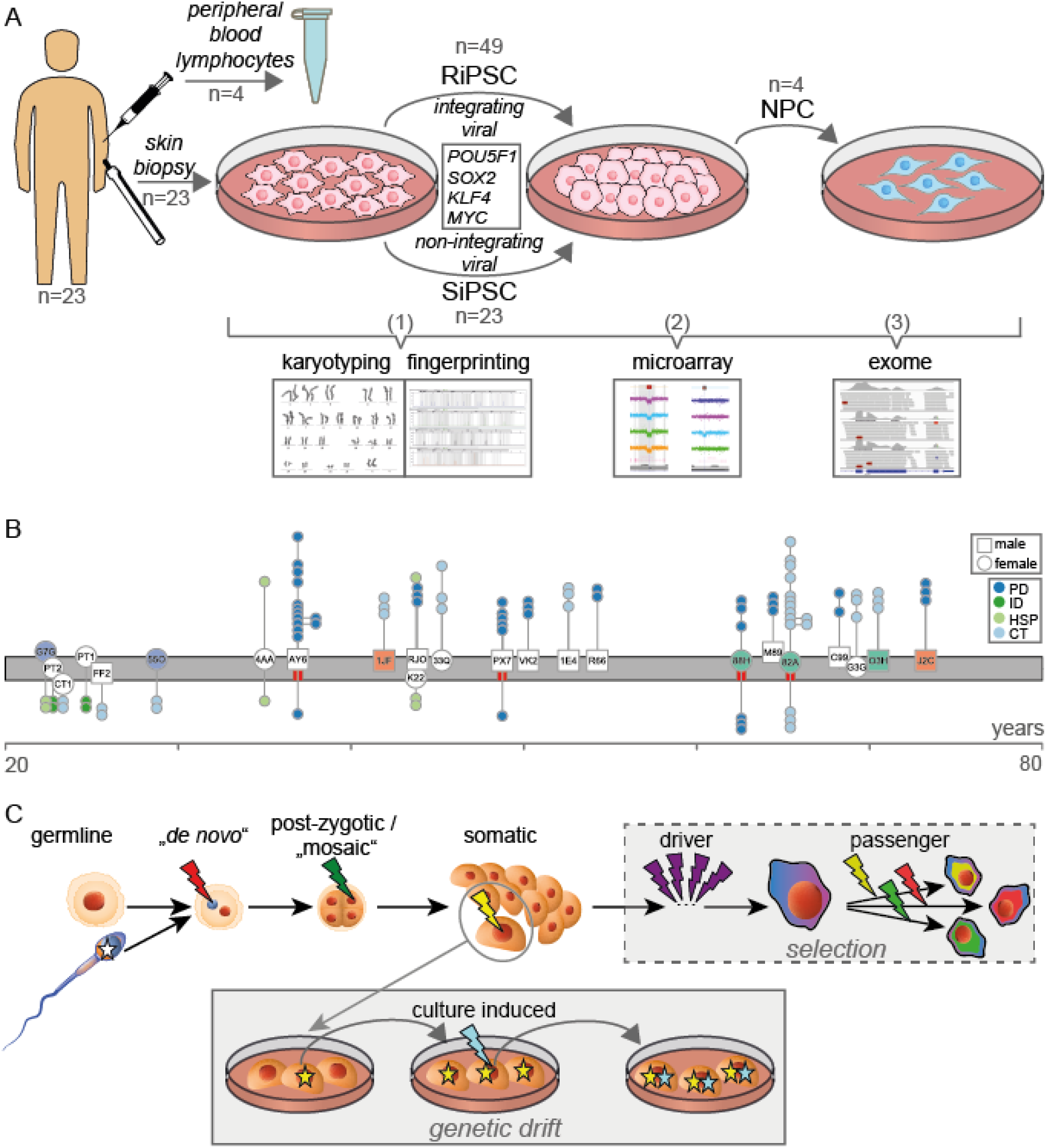
Schematic summary of the study and nomenclature of genetic aberrations. **(A)** Culturing and QC steps. Step 1: genetic fingerprinting and conventional karyotyping. Step 2: high-resolution CMA. Step 3: exome sequencing. **(B)** Graph showing the age distribution (x-axis) and phenotype of all donors. Fibroblast cultures are plotted as white symbols on the grey timeline (male = square; female = circle). The passage of the derived RiPSC (above) and SiPSC (below) cultures are plotted as circles connected to the respective fibroblast (y-axis; scattered for visualization). Derived NPCs are connected to the RiPSC they originated from. Red bars below the fibroblast symbols mark individuals with PBLs available selected for exome sequencing. See also File S1 for additional information. **(C)** Standardized nomenclature for variants/aberrations depending on the cell they arose in. The scheme compares the evolutionary history of a cancer cell (box “selection”) which is subject to a strong selective pressure with that of a cultured cell (box “genetic drift”) which is mainly subject to random genetic drift.

In the iPSC lines derived from fibroblasts, pluripotency was confirmed by positive staining for POU5F1 and NANOG for all iPSC lines, and fluorescence-activated cell scanning (FACS) analysis for TRA-1 −60 was positive for >90% of the cells in each line (Figure S1). All fibroblasts and iPSC lines generated in the ForIPS consortium, which passed these pluripotency criteria were send to genetic QC. A cell suspension from each culture was subject to an initial integrity screening (Figure 1A: “step 1”) using conventional karyotyping to detect aneuploidies and larger chromosomal aberrations. In this first QC step ~15%of iPSC cultures were discarded due to significant chromosomal aberrations (Figure S1 and File S2). In addition, DNA-based fingerprinting (PowerPlex assay) was employed to verify sample identity in most samples or was replaced by CMA based fingerprinting (Figure S1; File S2). Three iPSC lines did not match DNA from donor fibroblasts and were excluded from further analysis. For the remaining lines, fingerprinting matched with the respective fibroblast and with the reported donor sex. Samples which passed the first QC step were included into our subsequent studies (Figure 1B; File S1). This group included 72 primary iPSC lines with a median number of 3 iPSC lines per individual (range 2 to 6) and a median passage number of 14 at time of analysis (range 2 to 39). Forty-nine of these iPSC lines were generated by using integrating retroviral reprogramming (RiPSC) and 23 lines using non-integrating Sendai reprogramming (SiPSC) Yamanaka transcription factors.^2,16^ RiPSC had a higher median passage number of 15 (range 2 to 39) at analysis compared to 5 for SiPSCs (range 3 to 15). Four RiPSC lines from two individuals (“AY6”, “82A”) were differentiated into midbrain neuronal progenitor cells^1^(NPCs) and had a median passage number of 7.5 (range 5 to 13). To investigate the relationship between passage number and somatic variants, four RiPSC lines from the same individuals were cultured to higher passages of 30 and 40, respectively.

### Detection of somatic CNVs by high density SNP-based CMA

In a second analysis step all study samples passing step 1 (23 fibroblast cultures, 49 RiPSCs, 4 hereof derived NPCs, 4 RiPSCs at passages 30 and 40, and 23 SiPSCs) were screened for CNVs with a high-resolution, single-nucleotide polymorphism (SNP)-based chromosomal microarray (CMA). We used the Affymetrix CytoScan HD array as it is an established and reliable tool in routine germline diagnostics at our Center for Rare Diseases.^17,18^ Array QC measures passed manufacturer recommended thresholds in 97.2% of analyzed samples (105/108). The CMA data for the other three samples were only marginally below these thresholds and after manual review considered to be of sufficiently good quality (File S2; Figure S3). The CMA for each analyzed culture was visually screened by a trained expert (M.K.) for aberrations d100 kilobases (kb) and absent from donor fibroblasts (Supplementary information). We identified a total of 93 sub-chromosomal CNVs with sizes ranging from 100 kb to 6.4 Mb (megabases) including 48 deletions and 45 duplications (Figure 2). Most aberrations (91/93) were smaller than the lower detection limit of 5 to 10 Mb typically assumed for G-banded karyotyping.^19^In addition, we observed trisomy of chromosome 12 in three RiPSC cultures (“Ì1JF-R1-002”, “Í1E4-R1-012”, “Ì1E4-R1-016”), twice only present in a sub-population of cells. In the SiPSC line “CT1-S1-010” we detected a copy number gain affecting all terminal markers on chromosome 17q. Despite its size of 5.9 Mb this CNV was not detectable by conventional karyotyping. The chromosomal position indicated the possibility of an unbalanced translocation which was confirmed by fluorescence in situ hybridization (FISH) analysis as a 14p/17q unbalanced translocation probably of somatic origin (Figure 3A, B). While karyotyping and CNV analysis based on intensity data of chromosome 9 showed unremarkable results in SiPSC line “i82A-S1-004”, SNP allele peak distribution uncovered a copy neutral allelic imbalance on the long arm of chromosome 9 indicating a ~30% sub-clonal cell population carrying a partial uniparental isodisomy (Figure 3C; Figure S2).

**Figure 2.**
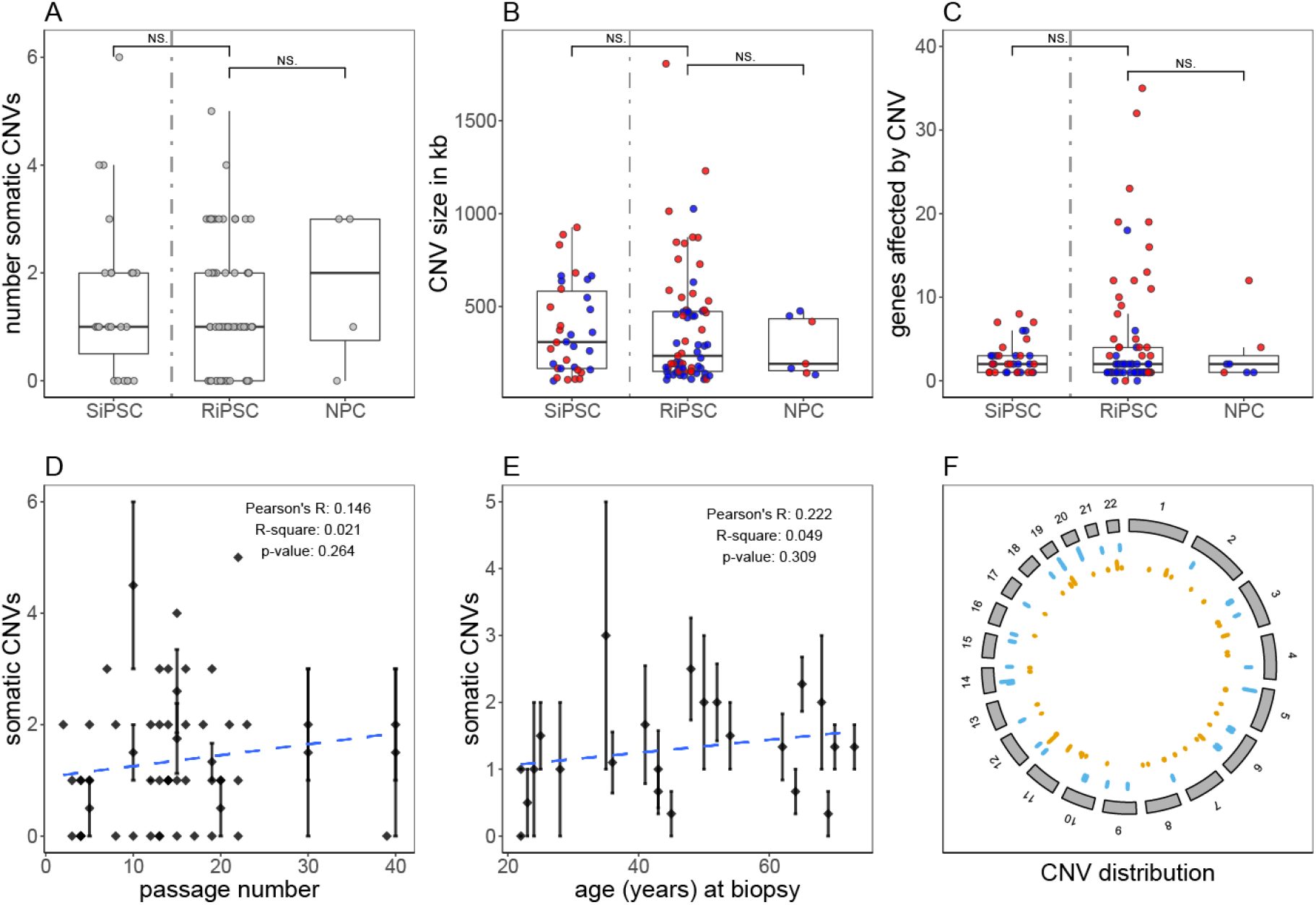
Summary of somatic CNVs identified in iPSC cultures by chromosomal microarray analysis (CMA). Box- and scatterplots for **(A)** the total number of somatic CNVs detected per analyzed cell culture sample (grey dots), **(B)** the genomic length (hg19) in kb of all detected somatic CNVs **(C)** and the number of affected genes (GenBank) within identified somatic CNVs (red dots = copy number loss, blue dots = copy number gain). SiPSC and RiPSC are separated by a grey dashed line. In the NPCs derived from the RiPSCs no new somatic CNVs were identified. No significant differences regarding number, size and gene content of somatic CNVs between RiPSC (n=49) and SiPSC clones (n=23) were detected (two sided Wilcoxon signed-rank test). Aneuploidies are not included and CNV outliers (one in SiPSC and one in RiPSC) sized over 5000 kb are excluded from panels B and C. **(D)** The average number of CNVs in all iPSCs grouped per individual and passage number plotted vs. the passage number. The dashed blue line represents the linear regression model fit (R^2^ = 0.021, p-value = 0.264). **(E)** The average number of CNVs in all iPSC grouped per individual plotted vs. the donor age in years at biopsy. The dashed blue line represents the linear regression model fit (R^2^ = 0.049, p-value = 0.309). Diamonds in D and E mark the respective average CNV count and are intersected by a standard error bar where applicable. **(F)** Circos plot showing the genomic (hg19) distribution of somatic CNVs in RiPSC (orange) and SiPSC (blue) clones. NS, not significant.

Next, we compared RiPSCs and SiPSCs to reveal method-specific differences: 58 somatic CNVs were detected in 34 of 49 (69.4%) RiPSCs, and 35 somatic CNVs in 17 of 23 (73.9%) SiPSCs. Only 15 of the RiPSCs (30.6%) and six SiPSCs (26.1%) showed no somatic CNVs. CNV size varied between 106 kb and 6.4 Mb in RiPSC, and between 100 kb and 5.9 Mb in SiPSC lines. The number of affected genes based on Genbank annotation varied between 0 and 139 with a higher variability in RiPSC lines. Three aberrations in RiPSC contained no genes, whereas all aberrations in SiPSC included genes. Our data showed no significant differences regarding number, size and gene content of somatic CNVs between RiPSC and SiPSC clones, indicating a comparable genetic cell quality (Figure 2A, B, C). Also, there was no significant difference between sexes, relatives- and affected-status confounding the analyses (Figure S3).

In four RiPSC clones cultured to higher passages we could not observe any CNV differences during passaging (File S3), and the average somatic CNV number aggregated per individual showed no correlation with passage number (Figure 2D). Additionally, the somatic CNV count was not correlating with the probands’ age at the time of biopsy (Figure 2E).

NPCs showed the same CNVs detected in the corresponding RiPSC clones indicating genetic stability during differentiation (Figure 2A, B, C; File S3). In the NPC culture derived from the RiPSC “i82A-R1-001” we observed two previously fixed CNVs which had lower intensities in the NPC compatible with a ~50% sub-population: A somatic deletion affecting the *DLG2* gene and a deletion affecting the genes *VCX* and *PNPLA4*(Figure S2). This observation shows that the RiPSC culture was initially oligoclonal, and points to either selective pressure of culture conditions or random genetic drift introduced by manual picking as the cause of the allelic shift in this NPC culture.

Although the identified somatic CNVs were scattered throughout the genome (Figure 2F), we detected three regions representing possible, specific hotspots. First, two overlapping deletions affecting the *CTNNA3* gene in 10q21.3 were identified in a RiPSC clone of individuals “88H” and “O3H”, respectively (Figure 3D, File S3). Second, three aberrations within the *DLG2* gene were detected: two overlapping deletions in the iPSC clones “i82A-R1-001” and “i82A-R1-002” of “82A” as well as a duplication in the SiPSC clone “iK22-S1-001” of “K22” (Figure S2). Many smaller and overlapping aberrations in both regions were observed in healthy control individuals (Database of Genomic Variants^20^). Furthermore, a mosaic gain in 20q11.21 including the *BCL2L1* gene was revealed in two different RiPSC clones of “PX7” and one clone of “1JF”.

**Figure 3.**
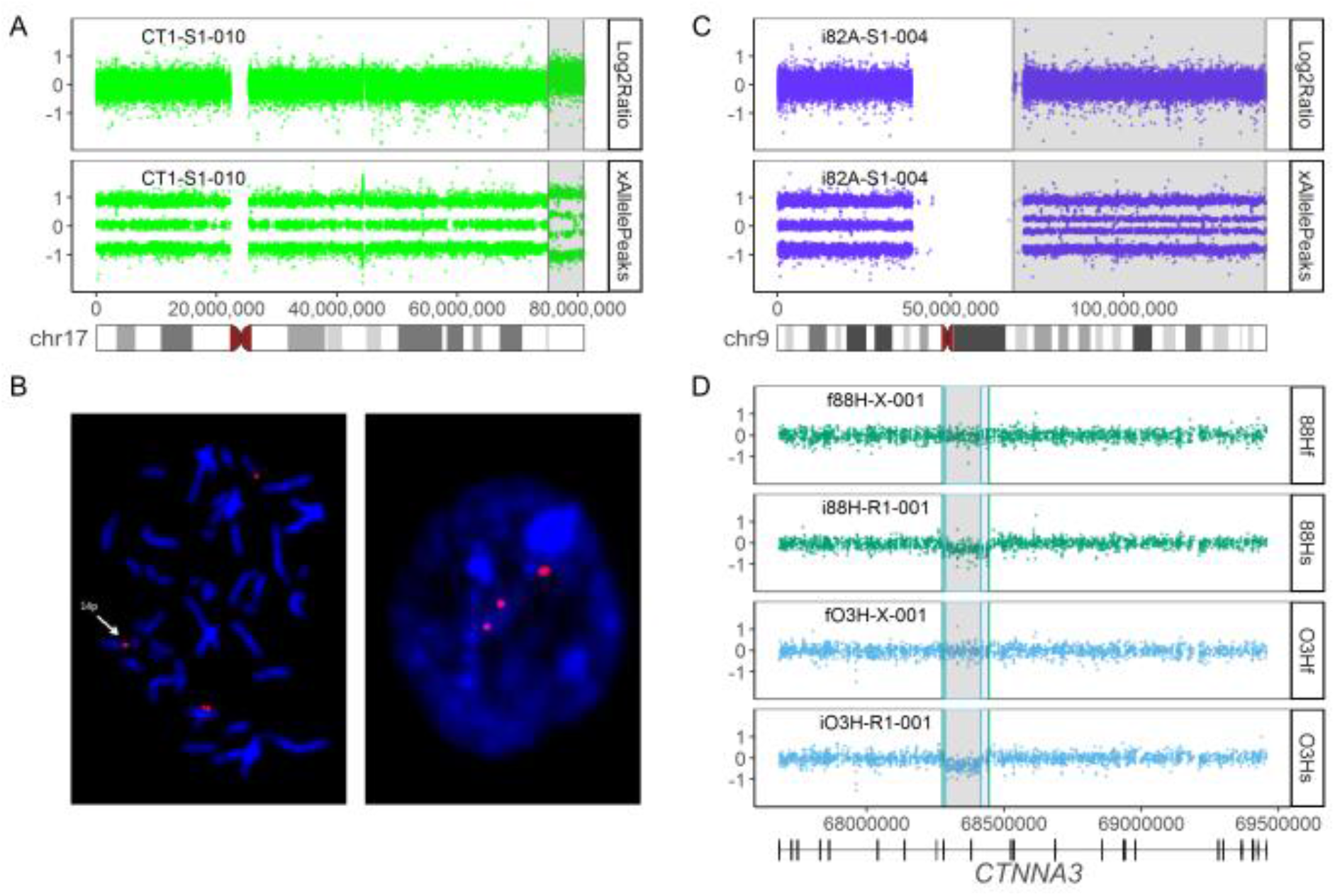
Examples of CNVs detected by SNP-based CMA. **(A)** Copy number analysis identified a chromosome 17q terminal gain not detectable with conventional karyotyping in the SiPSC line “CT1-S1-010”. **(B)** FISH analysis showing the unbalanced translocation 14p/17q in this clone (left = metaphase, right = interphase). **(C)** Conventional karyotyping and copy number analysis of chromosome 9 of the SiPSC line “i82A-S1-004” revealed unremarkable results (Log2Ratio top), but SNP allele peak distribution (xAllelePeaks bottom) uncovered a copy neutral allelic imbalance on the long arm of chromosome 9 (4 bands) while the short arm (left) shows normal allelic distribution (3 bands) (see also Figure S2). **(D)** Two independent overlapping intragenic deletions in the *CTNNA3* gene detected in the RiPSC lines “i88H-R1 −001 “ (green bottom) and “iO3H-R1-001” (blue bottom) and absent from their fibroblast cultures “f88H-X-001” (green top) and “iO3H-X-001” (blue top).

### Exome sequencing comparing iPSC and germline donor material to detect SNVs/indels

We selected a subset of samples for comparative exome sequencing with following inclusion criteria: (1) Availability of a germline DNA sample of the donor (blood) which was not a direct progenitor of the cultured cells (fibroblasts). (2) Availability of SiPSC, RiPSC and differentiated NPC lines of the same donor. (3) Access to higher passage samples of the lines. (4) Different affected status, age and sex. As the individuals “AY6”, “PX7”, “88H” and “82A” met these criteria, we selected a total of 34 samples (4 blood, 4 fibroblast, 8 RiPSC, 4 RiSPC passage 30, 4 RiSPSC passage 40, 6 SiPSC, 4 NPC). Exome sequencing on an Illumina HiSeq2500 machine and standard preprocessing resulted in aligned BAM files (Supplementary information) with a median on-target coverage of 163x (range 117x to 264x) and e95% of the exome target being covered by at least 20 reads (File S2).

Based on an initial feasibility test run with six exomes (File S1 and File S4; Figure S4; Supplementary information) and previous experience from pooled^21^ and somatic variant calling,^22^ we used the freebayes software,^23^ which simultaneously calls all classes of small nucleotide variants (SNVs = single nucleotide variants, MNPs = multiple nucleotide polymorphisms, indels = small insertions/deletions; when not specifically stated we use the term SNV/indel for all classes of small variants). All 34 exome samples were called together with 53 in-house controls from the same machine runs with freebayes and resulting variants were annotated with SnpEff.^24^ From here on we describe somatic variants obtained after applying hard filters to exclude variants with read evidence in the blood samples (Supplementary information; File S4). We considered resulting variants with alternate allele fractions (AF) ≥ 30% as fixed somatic and variants with AF < 30% as low frequency somatic variants (File S4; Figure S4). We identified a median of 38 fixed (minimum 17, maximum 256) and 1651 low frequency (minimum 739, maximum 3988) somatic SNVs/indels per sample in the coding target regions. We only report the results for the fixed variants and did not perform orthogonal validation (e.g. deep amplicon sequencing or digital PCR) for the low frequency somatic variants (see Figure S4) as previously analyzed by others.^13^

In analogy to the CNV analysis, we investigated SiPSC, RiPSC and NPC exome data for reprogramming or differentiation specific effects. No significant differences were detected for somatic SNV/indel numbers between RiPSC and SiPSC clones or between RiPSC and their derived NPCs (Figure 4A, B). Notably, the variance was higher for RiPSC (Figure 4A), an effect resulting from specific cultures (compare Figure 5A, B) with a much higher variant load.

**Figure 4.**
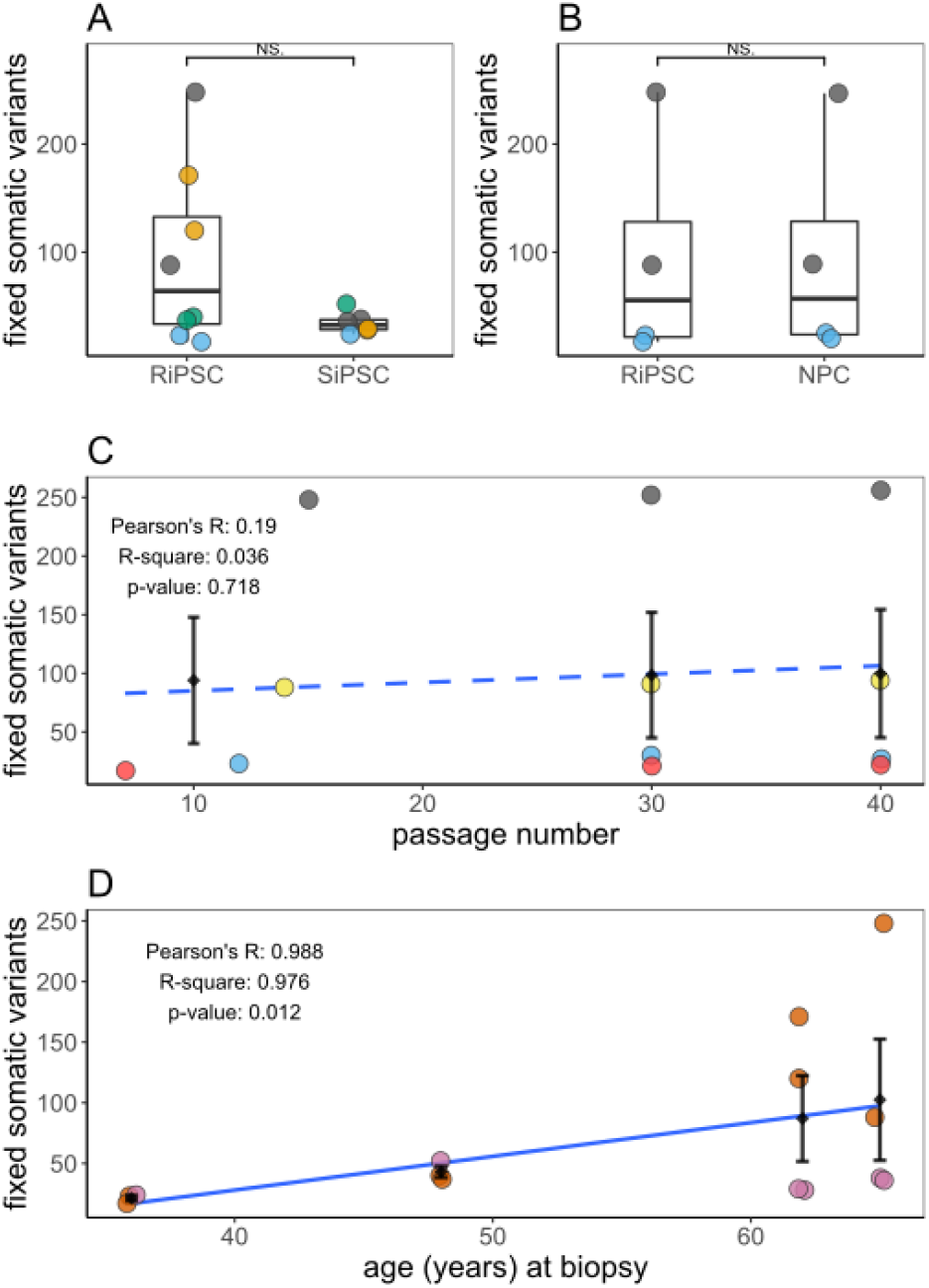
Summary of somatic SNVs/indels identified in iPSC cultures by exome sequencing. **(A)** Box- and scatterplot comparing the total number of fixed somatic SNVs/indels in independently reprogrammed SiPSC (n = 6) and RiPSCs (n = 8) from four donors (“82A”=grey, “88H”= orange, “AY6”= blue, “PX7”= green). **(B)** Box- and scatterplot comparing the total number of fixed somatic variants in RiPSC and derived NPCs from donors “82A” (grey) and “AY6” (blue). No significant differences were detected neither for somatic SNV/indel numbers between RiPSC and SiPSC clones nor between RiPSC and their derived NPCs (two sided Wilcoxon signed-rank test). Certain cultures have a much higher variant load (“82A”= grey, “88H”= orange). NPCs have the same variant profile as their progenitor cells. **(C)** Number of variants in four RiPSC lines (“i82A-R1-002”= grey, “i82A-R1-001”= yellow, “iAY6-R1-003”= blue, “iAY6-R1-004”= red) from donors “82A” and “AY6” cultured to higher passages vs. passage number. Diamonds mark the respective average SNV/indel count grouped by cell culture passage number (low passage numbers between 7 and 15 are considered as one group) intersected by a standard error bar. Dashed blue line represents the linear regression model fit using the actual passage number of the cells in the low group and the average of passage 30 and 40 (R^2^ = 0.036, p-value = 0.718). Note again the high spread influenced by the two cultures from individual “82A”. **(D)** The number of variants in all iPSC lines (RiPSC = ocher and SiPSC = lilac) from the four donors (n = 4 for “82A” and “88H”, n = 3 for “AY6” and “PX7”) plotted vs. the donor age. Diamonds mark the respective average SNV/indel count grouped by donor intersected by a standard error bar. Dashed blue line represents the linear regression model fit (R^2^ = 0.976, p-value = 0.012). NS, not significant.

Like for CNVs, we found no correlation between somatic SNV/indel variant load and passage number (Figure 4C). In contrast to the CNV analysis, the somatic SNV/indel count aggregated per individual showed a strong positive correlation with the probands’ age at the time of biopsy (Figure 4D). However, this observation is influenced by above mentioned iPSC cultures from older (Figure 5).

**Figure 5.**
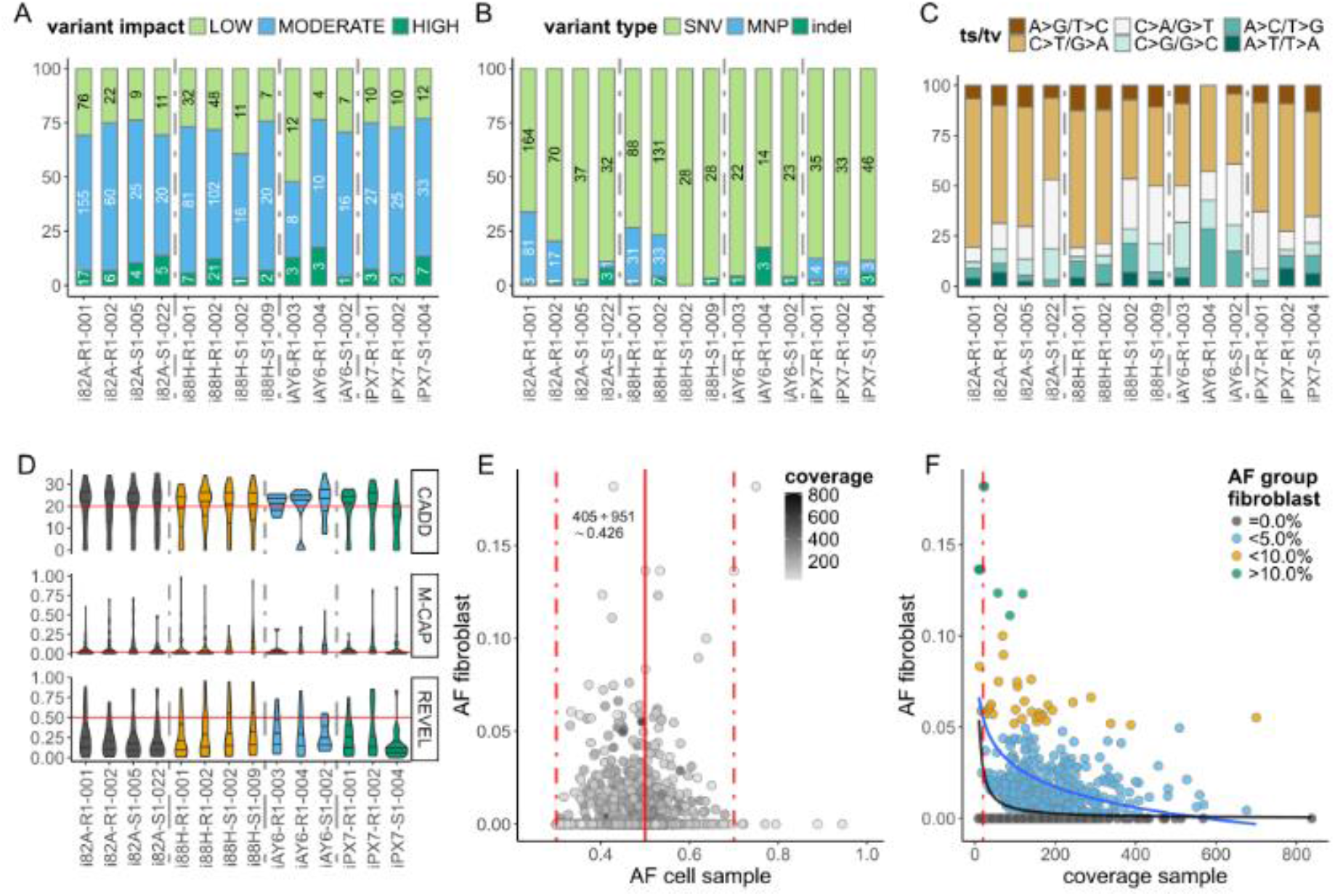
Mutational characteristics of somatic variants identified in iPSC cultures by exome sequencing. Stacked bar chart for the 14 primary RiPSC and SiPSC cultures from 4 individuals with passage numbers between 7 and 15 showing the relative number of variants partitioned **(A)** using SnpEff software annotated by variant impact group (HIGH = green, MODERATE = blue, LOW = light green), **(B)** by variant type (SNV= light green, MNP = blue, indel = green) and **(C)** by mutational subtype (transitions in brownish, transversions in greyish turquoise) of the SNVs in each iPSC sample. For A and B absolute variant counts are in the bars. **(D)** Distribution of three different SNV classifier scores represented as violin plots with median and quartiles. Red line represents the respective cutoff values (CADD = 20, M-CAP = 0.025, REVEL = 0.5). **(E)** Dot-plot showing the distribution of allele fraction (AF) in the analyzed iPSC cell cultures (x-axis) and their corresponding fibroblast culture (y-axis) with each point representing a variant shaded by read coverage in the iPSC exome (bright = low, dark = high read coverage at the respective variant position). Dotted vertical lines mark the expected AF for a heterozygous fixed variant (0.5) and typical variabilities seen in short read sequencing (0.3 to 0.7). **(F)** Dot-plot showing the relation between read coverage in the analyzed iPSC cultures and AF in the corresponding fibroblast culture. Dots are grouped and colored by fibroblast AF (no evidence in fibroblast = grey, d5.0%= blue, d10.0%= orange, >10%= green). The blue line represents the linear regression model fit (formula y ~ log(x); R^2^ = 0.202, p-value < 2.2e-16). The black line represents the theoretical AF in the fibroblast culture which is detectable at the respective coverage with a probability of 0.426 (variants with no evidence in fibroblast = 546, variants with at least 1 read in fibroblast = 405; 405/(405+546) ≈ 0.426) under a simple binomial draw model where one read is considered as sufficient evidence in the fibroblast. The red dotted line marks read coverage of below 20 where a high sampling variance is expected.

Next, we analyzed specific properties of the identified somatic SNVs/indels. Variants predicted to have a moderate impact on gene function (mainly missense variants) represent the largest proportion of identified somatic variants (range 35% to 69%) per sample (Figure 5A). In most iPSC samples, somatic variants were mainly SNVs, with only a small portion of indels and MNPs identified. However, four samples showed an unusual high proportion of MNPs (Figure 5B). A closer examination of these samples (File S4) showed that the MNPs are mainly CC>TT dinucleotide mutations at dipyrimidines and that they additionally had an increase in C>T/G>A transitions (Figure 5C), both mutational signatures typical for ultraviolet light (UV) irradiation damage.^25^

Missense variants represented a large part of the identified somatic SNVs in the iPSC cultures. Compared to truncating variants their functional interpretation is difficult. We used different computational prediction scores to assess their potential pathogenicity. Interestingly the scores obtained for a large portion (CADD: 44,1%, M-CAP: 35,0%, REVEL: 12,6%) of these somatic missense SNVs are above the respective recommended pathogenicity thresholds (Figure 5D).^26–28^

Our exome study design with concurrent sequencing and analysis of blood germline and parental fibroblast culture samples enabled us to search for evidence of low frequency somatic variants in fibroblasts due to polyclonality (“somatic mosaicism”). While low frequency variants in bulk sequencing data are inherently noisy when analyzed alone, prior knowledge of a fixed variant in a descendent culture sample increases the locus specific probability of low frequency reads being *bona fide* somatic variants.^13,29^ Accordingly, the allele fraction (AF) for fixed variants in the analyzed iPSC cell cultures followed an expected normal distribution of around 0.5, while most of the variants with read evidence in the fibroblasts had a lower AF. In addition, variants at the lower coverage tails had a larger variance in AF influenced by random sampling (Figure 5E; Figure S4). We found a correlation between read coverage at somatic variant positions in the iPSC cultures and AF in the corresponding fibroblast culture, indicating that somatic variants at low AF can only be found in the fibroblast if sufficient read coverage is available. Using a simple binomial draw model, we demonstrate that most variants potentially identifiable as being present in the fibroblasts (somatic) indeed do have reads supporting them (Figure 5F). It is likely that the remaining somatic variants are still somatic but only present at a very low AF in the original fibroblast culture and that they were just not detectable by bulk exome sequencing^13^.

### Multiple secondary analyses revealed additional iPSC culture characteristics

While the mitochondrial genome (“chrM”) is not targeted in most commercial exome designs, exome data still contain considerable mitochondrial coverage due to their high copy number in each cell. We calculated the average coverage of chrM (median 263x, minimum 66x, maximum 765x) and normalized it to the coverage of chromosome 1 (File S5). Fibroblast and R/SiPSC cell cultures showed a significantly higher mitochondrial genome dosage than NPC cultures and peripheral blood lymphocytes (PBLs) (Figure 6A).

**Figure 6.**
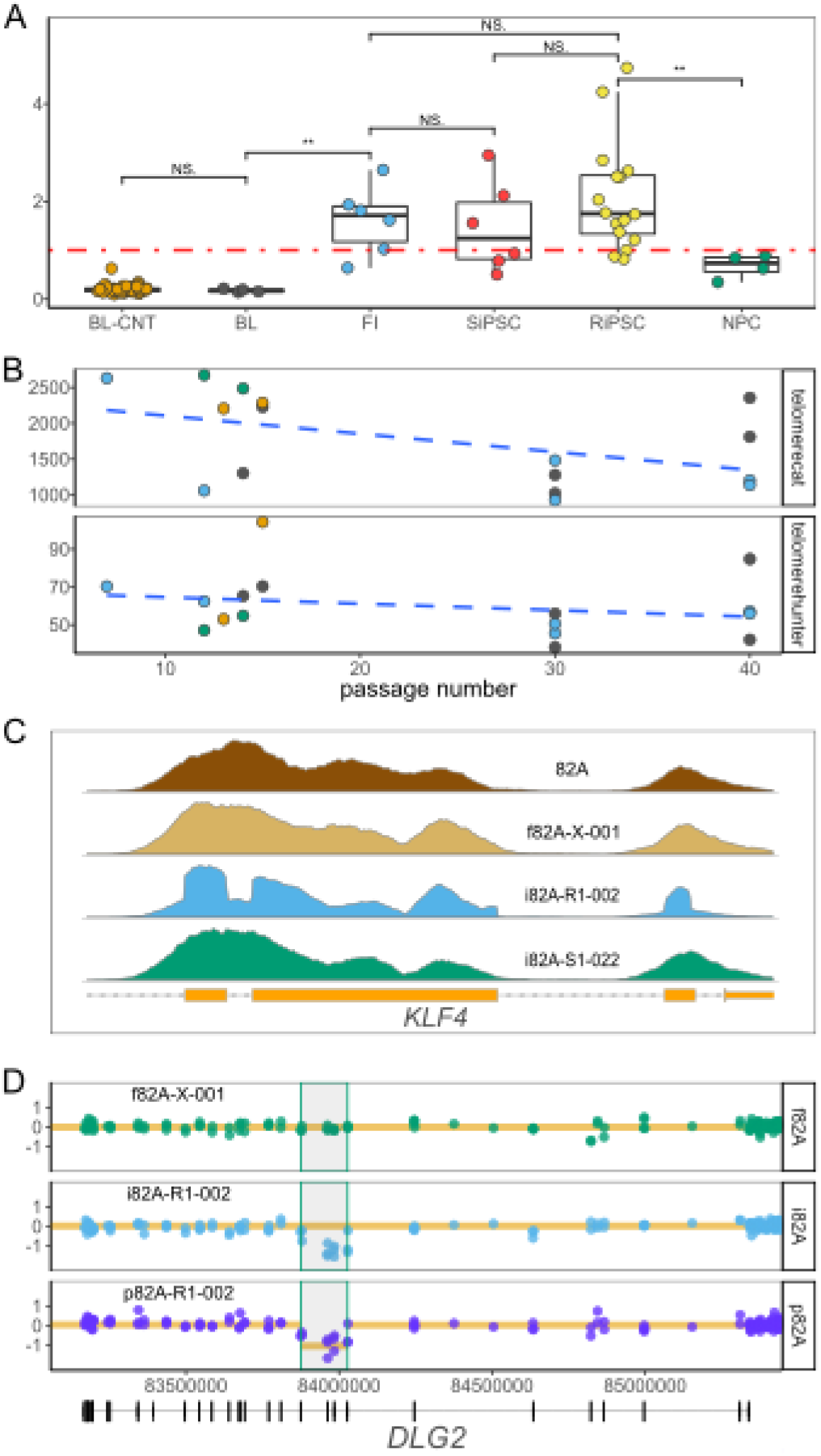
Exome sequencing enables multiple cellular analyses. **(A)** Box- and scatterplots of the relative mitochondrial genome ratio for all samples. Average read coverage for the mitochondrial genome (chrM) was normalized to the targeted regions of chromosome 1 (chr1). The level of significance is annotated by asterisks or as not significant (NS) (two sided Wilcoxon signed-rank test). Fibroblast (FI) and RiPSC/SiPSC cultures show a higher mitochondrial genome dosage than PBLs (BL = blood samples from individuals in this study; BL-CNT = blood samples from 53 inhouse control samples) and compared to NPC cultures. **(B)** Telomere content of all 16 RiPSC samples from the 4 individuals estimated from off-target telomeric reads by two different algorithms, telomerecat (upper panel) and telomerehunter (lower panel) plotted vs. the passage number. While both plots show a negative correlation of telomere content with higher passage number (telomerecat: Pearson’s r = −0.483, R^2^ = 0.233, p-value = 0.058; telomerehunter: Pearson’s r = −0.251, R^2^ = 0.062, p-value = 0.349) the results are not significant (see also Figure S5). **(C)**Comparison of the read coverage profile at the *KLF4* gene locus of different materials from individual “82A” (blood = brown, fibroblast = tan, SiPSC = green, RiPSC = blue). The sudden breaks at the exon-intron boundary indicate multiple integrations of a plasmid with a *KLF4* transcription factor insert which has no introns (see also Figure S6). **(D)** Example of a somatic deletion in the *DLG2* gene called from the exome data of the NPC sample (“p82A-R1-002”= dark blue) and absent in the corresponding fibroblast culture (“f82A-X-001”= green). Dots represent target or anti-target coverage bins (y-axis = log2 ratio) and the orange line marks the copy number call by the CNVkit algorithm^32^ for each segment. Note that the deletion was only called in the NPC and not in the RiPSC (“i82A-R1-002”= light blue) although the deletion had been previously confirmed in both samples by CMA (see also Figure S6). NS, not significant; “***”, 0.001; “**”, 0.01, “*”, 0.05.

Likewise, telomeric genomic regions are not targeted in exome designs but have a high relative coverage in the genome. We used two recently described software algorithms (telomerecat^30^, telomerehunter^31^) to compute the relative telomere content from exome data and to correlate it with the passage numbers. While the estimates from both algorithms showed a trend towards less telomere content in higher passages, these results were not significant (Figure 6B). It should be noted that the telomeric content of the 53 in-house exome controls used, when correlated with age, also showed a non-significant trend (Figure S5).

In our initial exome variant calling test in RiPSCs we identified variants in the *POU5F1* gene locus absent from the parental fibroblast. These were confirmed to be single nucleotide variants from the integrated viral vector (Figure S6). We therefore excluded the genomic regions of all transcription factors used for reprogramming from variant calling (Supplementary information). When examining these regions, we noticed the coverage profile of the RiPSCs having sudden breaks at the exon-intron boundaries like the profile seen in RNAseq. In contrast, fibroblasts and SiPSCs show bell-like shapes over the capture probes, which is typical for capture-based enrichment (Figure 6C). Our observation indicated multiple genomic integrations (Figure S6) of the plasmid with intron-free transcription factor inserts used for reprogramming of the RiPSC lines.

We wondered whether algorithms for CNV detection from exome data could replace or supplement the widely accepted CMA analysis. The CNVkit algorithm^32^ uses intergenic reads to achieve a more uniform marker coverage across the genome. While several CNVs detected previously by CMA were also called from exome data using this software, several others were missed (Figure 6D; Figure S6; File S3).

Off-target reads can also be used to check sequencing data for DNA of microorganisms like mycoplasma or cross-individual contamination. We used the MinHash based BBSketch algorithm (https://jgi.doe.gov/data-and-tools/bbtools/) to screen our exome files for cell culture contamination but did not find any evidence for high-grade contamination (Figure S5; File S5). Similarly, we could exclude significant cross-individual contamination, a known problem in iPSC cultures.^33^ using the ContEst^34^ software (Figure S5; File S5).

## DISCUSSION

Since the discovery of reprogramming methods for somatic cells into pluripotency, the stem cell field has rapidly progressed.^23^ Precise disease modelling and personalized treatment are some of the promises the iPSC technology is beginning to fulfill.^7^ Though advances are increasingly encouraging, there is still considerable heterogeneity in research practices.^4,35^ This is especially evident in genetic QC, which in recent years only has received systematic attention in large cohorts.^5,9^ Despite a wealth of available experience from pioneering genetic fields regarding rare diseases or cancer genetics, the community has not yet agreed upon common minimal standards for an iPSC line to be acceptable as a model and to be safe for therapeutic use. Here, we describe the application of diagnostic grade technologies to ensure genetic integrity for a collection of iPSCs and differentiated progeny cells from the ForIPS consortium.

We confirm the minimal standard of conventional karyotyping and genetic fingerprinting. G-banded karyotyping led to the exclusion of an appreciable proportion of cell lines with numerical chromosomal anomalies, at a comparable frequency with other reports.^36^ but also large structural chromosomal rearrangements, which are quite frequent in iPSCs (Figure 3A, B; Figure S1; File S2). While this technique is considered relatively cheap, it requires a lot of hands-on work and does not produce results in a computable electronic form. CMA analysis for copy number aberrations can also identify aneuploidies. However, chromosomal rearrangements in a balanced state would be missed (Figure 3C; Figure S1). Some groups perform optical mapping as an alternative screening method.^14^ Despite its currently higher costs and the need for specific DNA extraction methods, its higher resolution and computational accessibility might make optical mapping a method of choice for structural aberrations. Also, genetic fingerprinting proved to be a valuable first line QC step which allowed us to resolve sample mix-ups. While short tandem repeat (STR)-based methods, like the one we used, are widely employed for identity testing, these do not allow sample tracking in a complete genetic pipeline. A single nucleotide polymorphism based profiling panel for sample tracking^37^ would likely be more valuable for biobanks.

Our results using high density CMA showed that about 70%of iPSC lines have a detectable somatic CNV e100 kb, independent of the reprogramming method used (Figure 2A, B, C). This fraction is higher than in previous large reports,^5,9,38^ which can be attributed to variable CMA resolution and differences in filtering and analysis between the studies. Indeed, a smaller study using the same CMA platform we choose, did also find CNVs in a relatively large portion of iPSCs,^39^ We could point out several genomic regions affected by recurrent CNVs in iPSCs, explainable either by genetic fragility of the locus^40^or by proliferative survival advantage.^41^ By applying high coverage exome sequencing we identified SNV/indel variants in the coding gene regions in every iPSC line analyzed, independent of the reprogramming method used (Figure 4A). Interestingly, every primary iPSC line had at least one fixed somatic high impact (truncating) SNV/indel and several somatic missense variants of which a large portion was predicted as damaging to the protein function by different computational scores (Figure 5A, B, D). Several of the identified somatic variants affect genes implicated in cancer or monogenic diseases as well as genes with elevated expression in the brain (Table 1). These findings are well in line with previous reports.^12^ Our results suggest a functional impact of certain somatic variants in the iPSC lines. Together with the high variability in somatic variant load observed for all variant classes (Figure 2A, Figure 4A), even in isogenic lines, these observations signify that each line must be individually assessed before use in downstream experiments or therapeutic applications. In addition, we found no significant differences between integrating and non-integrating reprogramming methods regarding somatic CNVs (Figure 2A, B) and SNV/indel (Figure 4A) counts, thus supporting a recent publication for SNVs/indels.^14^ This information is of special value to researchers working with established RiPSC lines.

**Table 1.** Fixed variants with predicted loss-of-function effect in known cancer associated genes, OMIM disease genes or genes highly expressed in the brain.

| Sample | Gene | HGVS | List | OMIM-G | OMIM-P | Phenotype | Inh. | pLI |
| --- | --- | --- | --- | --- | --- | --- | --- | --- |
| i82A-S1-022 | IL1RAPL1 | c.1372+1G>T, p.? | OMIM, HPA | *300206 | #300143 | Mental retardation, XLR 21/34 | XLR | 1.00 |
| p82A-R1-001 | ADAT3 | c.485G>A, p.(Trp162*) | OMIM | *615302 | #615286 | Mental retardation, AR 36 | AR | 0.00 |
| i82A-R1-001 | RP1 | c.3949C>T, p.(Gln1317*) | OMIM | *603937 | #180100 | Retinitis pigmentosa 1 | AR, AD | 0.00 |
| i82A-S1-022 | MMP20 | c.1381dupA, p.(Thr461Asnfs*5) | OMIM | *604629 | #612529 | Amelogenesis imperfecta, type IIA2 | AR | 0.00 |
| i82A-R1-001 | GPR162 | c.747_748delinsTT, p.(Arg250*) | HPA | na | na | na | na | 0.02 |
| i88H-R1-002 | BRAF | c.981-2A>G, p.? | CCS, OMIM | *164757 | #115150, #613707, #613706 | Cardiofaciocutaneous syndrome; LEOPARD syndrome 3; Noonan syndrome 7 | AD, AD, AD | 1.00 |
| i88H-R1-002 | TWIST2 | c.3G>T, p.? | OMIM | *607556 | #200110, #209885, #227260 | Ablepharon-macrostomia syndrome; Barber-Say syndrome; Focal facial dermal dysplasia 3, Settleis type | AD, AD, AR | 0.44 |
| i88H-R1-002 | PAM16 | c.285_288del, p.(Ser96Argfs*44) | OMIM | *614336 | #613320 | Spondylometaphyseal dysplasia, Megarbane-Dagher-Melike type | AR | 0.14 |
| i88H-R1-001 | CDT1 | c.352-1G>A, p.? | OMIM | *605525 | #613804 | Meier-Gorlin syndrome 4 | AR | 0.00 |
| i88H-R1-002 | LAMB1 | c.869_870del, p.(Val290Glyfs*13) | OMIM | *150240 | #615191 | Lissencephaly 5 | AR | 0.00 |
| i88H-R1-002 | MTM1 | c.1497del, p.(Trp499Cysfs*3) | OMIM | *300415 | #310400 | Myotubular myopathy, XLR | XLR | 1.00 |
| i88H-R1-001 | DOCK2 | c.3060_3072+6del, p.? | OMIM | *603122 | #616433 | Immunodeficiency 40 | AR | 1.00 |
| i88H-R1-002 | CYP46A1 | c.894_897del, p.(Phe299Serfs*16) | HPA | *604087 | na | na | na | 0.75 |
| i88H-R1-002 | MCF2 | c.541G>T, p.(Glu181*) | HPA | *311030 | na | na | na | 0.94 |
| iAY6-R1-003 | GRIK2 | c.723+1G>A, p.? | OMIM, HPA | *138244 | #611092 | Mental retardation, AR, 6 | AR | 0.99 |
| iAY6-R1-003 | SYNE2 | c.4051C>T, p.(Gln1351*) | OMIM | *608442 | #612999 | Emery-Dreifuss muscular dystrophy 5 | AD | 0.00 |
| iAY6-R1-003 | C2CD3 | c.1726_1730+2delinsC, p.? | OMIM | *615944 | #615948 | Orofaciodigital syndrome XIV | AR | 0.00 |
| iAY6-R1-003 | ANKRD11 | c.5759_5763delinsG, p.(Thr1920Argfs*42) | OMIM | *611192 | #148050 | KBG syndrome | AD | 1.00 |
| iPX7-R1-001 | CARD11 | c.214C>T, p.(Arg72*) | CCS, OMIM | *607210 | #616452, #615206, #617638 | B-cell expansion with NFKB and T-cell anergy; Immunodeficiency 11A; Immunodeficiency 11B | AD, AR, AD | 1.00 |
| iPX7-R1-001 | ALG2 | c.32C>A, p.(Ser11*) | OMIM | *607905 | #616228 | Myasthenic syndrome, congenital, 14, with tubular aggregates | AR | 0.02 |
| iPX7-S1-004 | DNAH5 | c.2049del, p.(Gln684Lysfs*7) | OMIM | *603335 | #608644 | Ciliary dyskinesia, primary, 3, with or without situs inversus | AR | 0.00 |
| iPX7-S1-004 | ITCH | c.540dup, p.(Cys181Leufs*7) | OMIM | *606409 | #613385 | Autoimmune disease, multisystem, with facial dysmorphism | AR | 1.00 |
| iPX7-S1-004 | VCAN | c.2492_2495del, p.(Leu831Glnfs*5) | OMIM | *118661 | #143200 | Wagner syndrome 1 | AD | 1.00 |
| iPX7-S1-004 | ARPP21 | c.1923T>G, p.(Tyr641*) | HPA | *605488 | na | na | na | 0.00 |
| iPX7-S1-004 | WBSCR17 | c.1081-1_1081delinsAA, p.? | HPA | *615137 | na | na | na | 0.10 |

Inh., inheritance mode; HGVS, Human Genome Variation Society nomenclature; OMIM-G, OMIM gene number, OMIM-P, OMIM phenotype number, pLI, probability of loss-of-function intolerance<sup>56</sup>.

The relationship between culture passaging and somatic variants count has been controversially discussed in the literature. While early analyses have described a negative correlation between CNV count and passaging,^42^ recent studies using low resolution CMA^5,12^ or whole genome sequencing^13^ could not confirm this. Furthermore, an older study showed an increase in coding SNV counts from 7 to 13 for a single analyzed iPSC line between passage 9 and 40,^43^ Our results do not support a strong effect of passaging on either CNV or SNV/indel counts (Figure 2E, Figure 4C). The four NPC lines differentiated from RiPSCs in our study showed no additional CNVs (Figure 2A, B, C) and have not significantly acquired SNVs/indels during differentiation (Figure 4B). Together, these data argue against a strong effect of passage number on somatic variant count.

Based on increasing numbers of somatic CNVs in aging individuals as demonstrated in cancer studies,^44–46^ one would expect to find higher frequencies of this mutation type in iPSCs derived from older donors. Our results, however, demonstrated no significant correlation between donor age and somatic CNV count, confirming similar recent reports.^5,12^ In contrast to CNVs, somatic SNV/indel load in exome regions has been shown to linearly increase with donor age in iPSCs derived from peripheral blood mononuclear cells.^12^ We also confirm this observation in our iPSC sample collection derived from skin fibroblasts (Figure 4D). Altogether, our findings and the descriptions in the literature point to differences in the mutational mechanisms and cellular processes involved in the formation of somatic CNVs and SNVs/indels. Our results point to UV irradiation damage related somatic sub-clonality in the parental fibroblasts as a source for SNVs/MNPs and inter-culture variability (Figure 4A, D; Figure 5A, B, C). Recent studies suggest that most variants identified in iPSC, but absent from the donor germline, are already present in a subpopulation of the cells of origin.^12,13,15^ We also show extensive somatic mosaicism in the parental fibroblast cultures as a source for fixed somatic variants in iPSCs (Figure 5F). Considering the data regarding passaging, we propose that random genetic drift induced by colony picking from poly/oligoclonal cell cultures and not positive selection is a major cause of somatic variation in iPSC clones (Figure 1C). This model is very different from the typical situation in cancer, where few “driver” mutations pose a strong advantage^47^ in an environment of selective pressure, while most “passenger” variants are neutral (Figure 1C). The goal in iPSC research is not to find detrimental driver mutations but to produce intact cells resembling the donor, thus successful strategies in cancer and iPSC fields will differ.

Mitochondria are crucial for cellular senescence and pluripotency in iPSCs^48^. Differences in mitochondrial morphology, count^49^ and mitochondrial DNA (mtDNA) content^50,51^ during pluripotent stem cell reprogramming and differentiation have been reported.^52^ Our analysis of the mitochondrial genome content showed significant differences between PBLs, iPSCs and differentiated NPCs, but not between fibroblasts and iPSCs (Figure 6A). A similar method for relative quantification of mtDNA from exome data has recently been compared to gold standard methods.^53^ These data highlight the added value of high-throughput sequencing reads for complementary analyses with potential use in iPSC characterization. The application of our method in large studies will likely expand our current knowledge of mitochondrial function in iPSCs and their progeny. Our exemplary attempts to telomere content analysis, viral integration and CNV analysis from exome data show that these analyses are in principal possible but need further evaluation and calibration (Figure 6B, C, D). Albeit applicable to exome data, most of the described techniques will likely lead to better results using whole genome sequencing data.

In conclusion, we applied high-resolution diagnostic methods in a systematic pipeline to ensure genetic stability of iPSCs generated in the ForIPS consortium and confirmed several previous associations in an iPSC collection from diverse donors. Most importantly, we showed that different clones have a high variability regarding somatic variant load. This highlights that the genetic evaluation of each individual iPSC clone is fundamental prior to its use as model or for therapeutic purposes. A combination of karyotyping by optical mapping, CMA and exome sequencing will likely provide the best combination regarding cost and efficiency in the next years. As even the smallest variant classes can have detrimental effects on important genes (Table 1), we recommend an inspection of all iPSCs based on three pillars: karyotyping for balanced aberrations, CMA for CNV detection, and NGS to search for SNVs/indels. Ideally these analyses should be performed on the initial iPSC cultures in comparison to an independent germline sample to find the best iPSC line before using these for experiments and again on later derivatives to ensure validity of functional results before publication.

## METHODS

### Inclusion of subjects in the ForIPS resource

The ForIPS research consortium (http://forips.med.fau.de/) has established an institutional iPSC biobank resource to explore diseases of the brain, particularly PD. All reported iPSC lines with adequate consent have been registered in hPSCreg^54^. To exchange selected lines for research purposes the scientific board of the UKER biobank will consider each request.

Twenty-three individuals were recruited at the Department of Molecular Neurology (Universitätsklinikum Erlangen). All individuals were phenotypically examined by a clinician experienced with neurological diseases. PD patients were diagnosed by board-examined movement disorder specialists according to consensus criteria of the German Society of Neurology, which are similar to the UK PD Society Brain Bank criteria for diagnosis of PD^55^. Age at tissue donation, gender, ethnicity and family history were assessed. All participants gave written informed consent to the study prior to donating a skin biopsy from a typically sun unexposed area of the inner upper arm. From this biopsy, a fibroblast stock culture was created. Four individuals additionally donated PBLs for an independent germline DNA sample (Figure 1A). Symptomatic individuals had targeted genetic testing to exclude or confirm monogenic forms of PD, HSP and ID (see Supplementary information). Study approval was granted by the local ethics committee (No. 4485 and 4120).

### Reprogramming, differentiation, culture conditions and genetic QC

Detailed methods used for generation of iPSC, differentiation of NPCs, cell culture conditions and for the genetic QC analyses performed are described in the Supplementary information.

## ACKNOWLEDGMENTS

We thank all participating individuals for donating materials. The authors thank Brigitte Dintenfelder and Michaela Kirsch for excellent technical assistance. This study was supported by the Bavarian Ministry of Education and Culture, Science and the Arts within the framework of the Bavarian Network for Induced Pluripotent Stem Cells: ForIPS. Additional support came from the Bavarian Molecular Biosystems Research Network: BioSysNet, the German Federal Ministry of Education and Research (BMBF: 01GQ113, 01GM1520A, 01EK1609B), the DFG funded research training group GRK2162 (B.W., M.R., J.W., A.R.), and the Interdisciplinary Centre for Clinical Research (University Hospital of Erlangen, E23 to A.R., E25 to B.W., J52 to M.R.).

## AUTHOR CONTRIBUTIONS

A.Re. and B.W. conceived and supervised the study. B.P., M.K., B.W. and A.Re. conceived the methodology. Z.K., J.W., R.A., A.Ra. and F.E. provided patient samples and clinical data. S.P., A.S., J.G, R.A. and M.F. generated iPSC and NPC cultures and verified the pluripotency of iPSCs. C.K. and M.K. performed DNA extraction and genetic fingerprinting. A.B.E. and S.U. generated data for molecular karyotyping and high-throughput sequencing. U.T. and M.K. performed karyotyping and FISH analysis. B.P. and M.K. analyzed and interpreted the molecular data. B.P. and M.K. prepared figures and tables. B.P., M.K., B.W. and A.Re. wrote and edited the manuscript with input from all co-authors.

## ADDITIONAL INFORMATION

The consent and ethics approval for the ForIPS study does not cover the deposition of identifiable germ line genomic data of study participants into public repositories. We follow the DFG (German Research Foundation) recommendations for safeguarding good scientific practice and thus internally archive all data for this study. We provide file checksums for all primary array and sequencing data (File S2). These shall be accessible for any legitimate request from the corresponding author (A.Re.). With future consent updates we plan to submit this genetic data to public repositories.

